# Methamphetamine induced regional-specific transcriptomic and epigenetic changes in the rat brain

**DOI:** 10.1101/2022.06.13.496004

**Authors:** Benpeng Miao, Xiaoyun Xing, Viktoriia Bazylianska, Pamela Madden, Anna Moszczynska, Bo Zhang

## Abstract

**Background:** Methamphetamine (METH) is a highly addictive central nervous system stimulant. Chronic use of METH is associated with multiple neurological and psychiatric disorders. An overdose of METH can cause brain damage and even death. Mounting evidence indicates that epigenetic changes and functional impairment in the brain occur due to addictive drug exposures. However, the responses of different brain regions to a METH overdose remain unclear.

**Results:** We investigated the transcriptomic and epigenetic responses to a METH overdose in four regions of the rat brain, including the nucleus accumbens, dentate gyrus, Ammon’s horn, and subventricular zone. We found that 24 hours after METH overdose, 15.6% of genes showed changes in expression and 27.6% of open chromatin regions exhibited altered chromatin accessibility in all four rat brain regions. Interestingly, only a few of those differentially expressed genes and differentially accessible regions were affected simultaneously. Among four rat brain regions analyzed, 149 transcription factors and 31 epigenetic factors were significantly affected by METH overdose. METH overdose also resulted in opposite-direction changes in regulation patterns of both gene and chromatin accessibility between the dentate gyrus and Ammon’s horn. Approximately 70% of chromatin-accessible regions with METH-induced alterations in the rat brain are conserved at the sequence level in the human genome, and they are highly enriched in neurological processes. Many of these conserved regions are active brain-specific enhancers and harbor SNPs associated with human neurological functions and diseases.

**Conclusion:** Our results indicate strong region-specific transcriptomic and epigenetic responses to a METH overdose in distinct rat brain regions. We describe the conservation of region-specific gene regulatory networks associated with METH overdose. Overall, our study provides clues toward a better understanding of the molecular responses to METH overdose in the human brain.

## Background

Methamphetamine (METH) is a powerful and widely abused psychostimulant that has deleterious effects on the central nervous system and results in addiction, which is a major public concern globally [1-4]. In the United States, close to 2,000,000 people use METH, and deaths from METH overdose are rapidly rising [5, 6]. Between 2015 and 2019, METH use increased by 43%, while the number of people suffering from METH use disorder increased by 62%. The number of deaths from METH overdose began to rise markedly increasing 10-fold between 2009 and 2019 [7]. METH use, particularly at high doses, is associated with neurologic and psychiatric disorders, as it causes severe cognitive impairment and neurobehavioral abnormalities [8, 9]. Recent studies have shown the acute and long-term effects of METH on cognitive functions such as attention, working memory, and learning, and METH overuse is often fatal [10-15]. Meanwhile, the high relapse frequency of METH abuse is a crucial challenge for treating the METH addiction [16-18]. There is no FDA-approved medication for METH use disorder, highlighting the importance of better understanding the molecular mechanism of the brain’s reaction to METH exposure and particularly to METH overdose, which causes neurotoxicity in the reward circuitry.

The reward circuitry plays a central role in different substance use disorders. The circuitry encompasses multiple brain subregions, including the ventral tegmental area, nucleus accumbens (NAc), dorsal striatum, amygdala, hippocampus, and regions of the prefrontal cortex [19-22]. In these different brain regions, distinct significant transcriptional and epigenetic changes can be caused by different addictive substances [19, 23, 24], creating complicated crosstalk between the epigenetic landscape and transcriptome in the brain [25, 26]. For example, acute or chronic exposure of the brain to psychostimulants, opiates, and alcohol can upregulate histone acetyltransferases (HATs) while suppressing histone deacetylases (HDACs), resulting in increased acetylation levels of histones H3 and H4 in the NAc and subsequent upregulation of their target genes [27-33]. METH exposure increased the acetylation levels of H4K5 and H4K8 in gene promoter regions in the rat striatum [34] and NAc [35]. Addictive drugs can also influence the expression of DNA methyltransferases (DNMTs) and further induce changes in DNA methylation, which plays essential roles in the cognitive learning and memory [36, 37]. Acute and chronic METH injections can increase DNMT1 expression in the rat NAc and dorsal striatum [38], and METH self-administration can increase the DNA methylation level of several potassium channel genes in the rat brain [39].

Recently, assay for transposase-accessible chromatin with high-throughput sequencing (ATAC-seq), a method for mapping genome-wide chromatin accessibility, has been widely used in addiction research to explore the open chromatin regions (OCRs) associated with exposure to addictive substances [40-44], including METH [45]. The changes in chromatin accessibility in OCRs interact directly with histone modifications and DNA methylation and are usually associated with the binding of different transcription factors (TFs), which are the regulatory hubs in the response to substance exposure. Many transcription factors, such as ΔFOSB, early growth response factors (EGRs), and multiple myocyte-specific enhancer factor 2 (MEF2), were found to respond to addictive substance stimuli and to regulate their downstream target genes [19, 23, 46-50]. However, most of these studies focused on a single brain region, most frequently the NAc. There is still a vast gap in understanding the simultaneous molecular changes in different brain regions upon substance stimuli.

In this study, we analyzed the dynamics of transcription and chromatin accessibility after METH overdose in four rat brain regions, including the NAc, dentate gyrus (DG), Ammon’s horn (CA), and subventricular zone (SVZ). The SVZ and the subgranular zone (SGZ) of the DG are the only neurogenic areas in the adult brain. Exposure to METH affects adult neurogenesis in the SVZ and SGZ [51, 52], while modulation of SGZ and SVZ neurogenesis impacts hippocampal-based cognitive function [53]. We determined that METH overdose induced a total of 2,254 differentially expressed genes (DEGs) and 25,598 differentially accessible regions (DARs) in the four rat brain regions. These four rat brain regions generally displayed a strong region-specific response to METH exposure at both the transcriptomic and epigenetic levels. Few of those DEGs and DARs were simultaneously affected by METH overdose in all four regions. We observed an interesting opposite regulation pattern between the CA and DG regions: 119 genes and 764 OCRs were significantly oppositely regulated by METH exposure.

Furthermore, METH overdose significantly affected the expression of 149 transcription factors (TFs) and 31 epigenetic modifiers (epigenes), which could regulate downstream gene expression and initiate epigenetic signatures. For accessible DARs identified in four regions, approximately 70% were orthologous in rat, mouse, and human genomes and were highly related to neurological processes. Some orthologous DARs in mouse and human genome regions could intersect with validated enhancers, and GWAS SNPs related to orthologous DARs in mouse and human genome regions could cross with validated enhancers and GWAS SNPs related to neuron biology. Meanwhile, TF binding motifs enriched in DARs composed distinct gene regulation networks in different brain regions in response to METH overdose. Taken together, our study provides a comprehensive multiomics investigation into the reactions of different brain regions exposed to METH overdose.

## Results

### RNA-seq and ATAC-seq data of 4 brain regions in normal and METH-overdosed rats

To explore the molecular changes in the brain in response to acute METH overdose, we exposed Sprague–Dawley rats to a high-dose METH binge (4 × 10 mg/kg, every two hours). At the same time, control group rats received saline injections. The NAc, DG, CA, and SVZ were collected 24 h after the last dose of METH or saline. The transcriptome and epigenome of the rats were assessed using RNA-seq and ATAC-seq assays (Fig. 1a and Additional file 1). As expected, we observed strong region-specific transcriptome and epigenome profiles in each of the four brain regions (Fig. 1b). Principal component analysis (PCA) separated the data from these four different rat brain regions in both RNA-seq and ATAC-seq assays (Fig. 1c), emphasizing the existence of strong brain-regional specificity at both the transcriptome and epigenetic levels. The DG and CA, two connected regions in the hippocampal formation, were closer to each other, as expected (Fig. 1c), but showed the opposite response to METH-overdose treatment, especially at the genome-wide chromatin level (Fig. 1c).

**Fig. 1.**
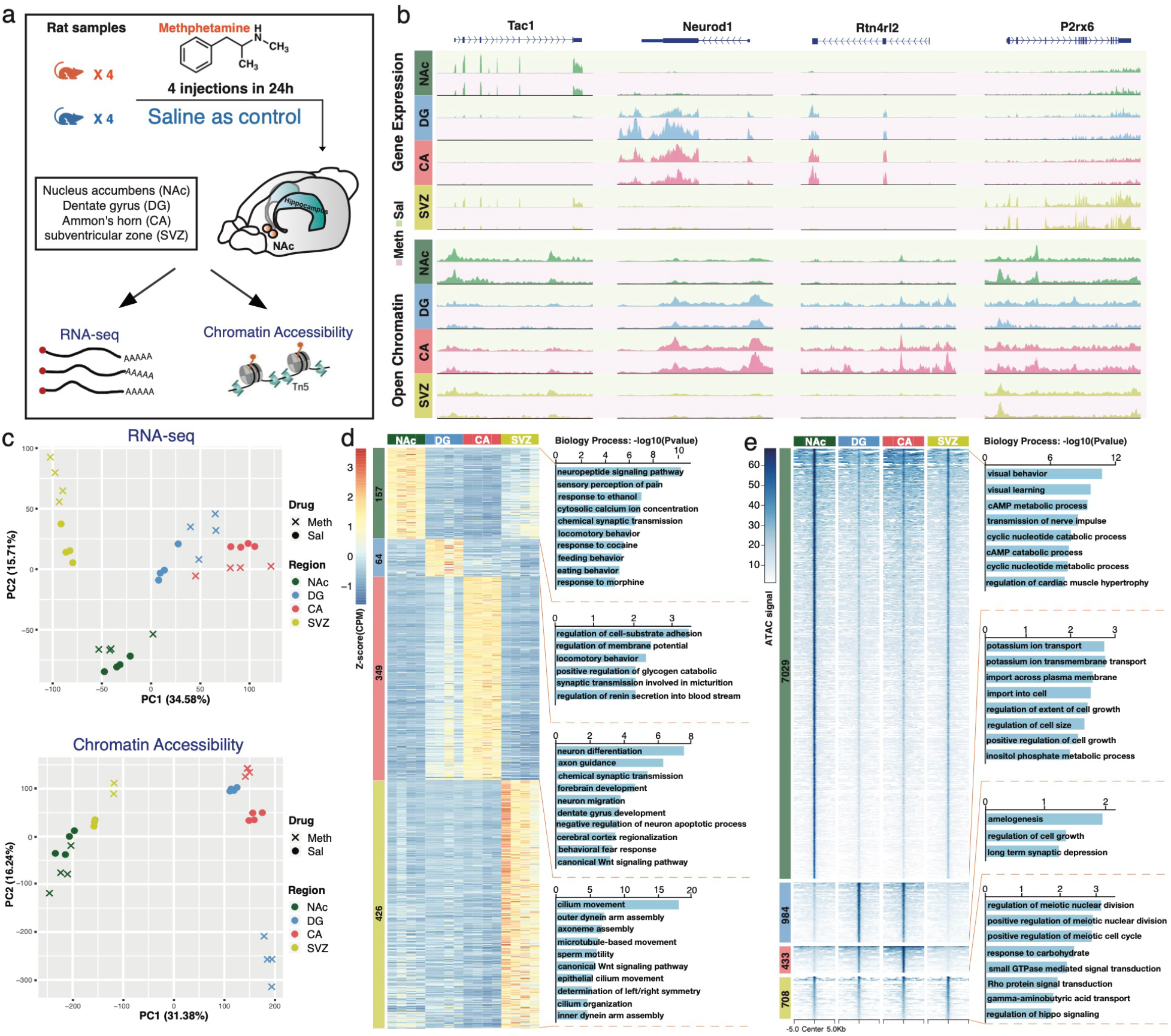
Gene expression and epigenetic signals of 4 brain regions from control and METH-overdose rat samples. **a**, Schematic outline of the experimental design. **b**, Examples of gene expression and open chromatin signals in METH-overdose and Saline samples of 4 rat brain regions. The *Tac1* gene was more highly expressed and became more open only in the NAc region. The *Neurod1* and *Rtn4rl2* genes showed high expression and chromatin accessibility in both the DG and CA. The *P2rx6* gene showed higher expression only in the SVZ. METH: METH-overdose samples, red background; Sal: saline samples, green background. **c**, The principal analysis (PCA) of METH-overdose and Sal samples of 4 brain regions for RNA-seq and ATAC-seq data. Cross: METH-overdose samples; dot: saline samples. The region-specific signatures of 4 brain regions in saline controls at both the transcriptomic (d) and epigenetic (e) levels. **d**, left part shows the number and expression Z score of region-specific genes in 4 brain regions; **e**, left part shows the number and open chromatin signals of region-specific accessible regions in 4 brain regions. The right parts of d and e show the enriched biological process terms.

We first investigated the region-specific signatures of the four brain regions in saline controls at both the transcriptomic and epigenetic levels by identifying the region-specific expressed genes and accessible chromatin regions (Additional file 2: Figure S1 and Additional file 3). At the gene level, 157, 64, 349, and 426 region-specific genes were identified separately in the NAc, DG, CA, and SVZ regions, respectively (Additional file 2: Figure S1e). These genes were significantly enriched in distinct neurological functions and processes in a region-specific fashion (Fig. 1d and Additional file 4). In the NAc region, 10 of 157 genes were directly associated with the drug responses to cocaine and morphine. CA region-specific genes were highly enriched in neuron differentiation and axon guidance. We identified 426 genes explicitly expressed in the SVZ region, and these genes were enriched in movement- and assembly-associated biological functions, suggesting their involvement in neuron differentiation and migration activities in the SVZ region (Fig. 1d). At the epigenetic level, we identified 7,029 NAc-specific open chromatin regions (OCRs, Additional file 2: Figure S1f). The genes around these NAc-specific OCRs were enriched in behavior, learning, and cAMP metabolic processes (Fig. 1e and Additional file 4). However, only a few hundred region-specific OCRs were identified separately in the other three brain areas (Fig. 1e and Additional file 2: Figure S1f). Such results suggested that the NAc brain area is unique at the epigenetic regulatory level.

### METH-overdose induced region-specific differentially expressed genes in 4 rat brain regions

METH overdose induced significant molecular changes in all four brain regions (Fig. 1c). To better characterize the region-specific transcriptomic response to METH overdose, we evaluated the differentially expressed genes in each region separately (Fig. 2a and Additional file 5). We identified 1,209 DEGs in the DG, the highest number among all four areas, indicating that the DG is the brain region that is most sensitive to METH overdose (Fig. 2a). In the SVZ, 957 genes were significantly changed, suggesting that the SVZ is a vital target brain region for METH-overdose stimuli. In the NAc and AC, only 358 and 329 genes, respectively, exhibited significant changes in expression. We further cross-referenced the METH overdose-responsive DEGs in all four brain regions and found that most of the DEGs were regionally regulated; only a few genes responded to METH overdose in multiple areas, including Gfap, Nrn1, and Drd1 (Fig. 2b and Additional file 2: Figure S2a). Such results suggested that the significant transcriptomic response to METH-overdose stimulus has high regional specificity and different sensitivity to stimulation in distinct brain regions. GO enrichment analysis of the region-specific DEGs revealed their enrichment in certain biological processes, e.g., specifically downregulated DEGs in the NAc, DG, and SVZ were enhanced in locomotor behavior and response to drugs, including amphetamine, morphine, and cocaine (Fig. 2c). METH overdose-induced upregulated DEGs in the NAc, DG, and CA were enriched in essential neuronal functions, such as signaling pathways, neuron development, axon guidance, memory, etc. Upregulated DEGs in the SVZ were highly enriched in myelination and oligodendrocyte development, suggesting that glial cells in the SVZ region have a distinct response to the METH-overdose stimulus (Fig. 2c and Additional file 6).

**Fig. 2.**
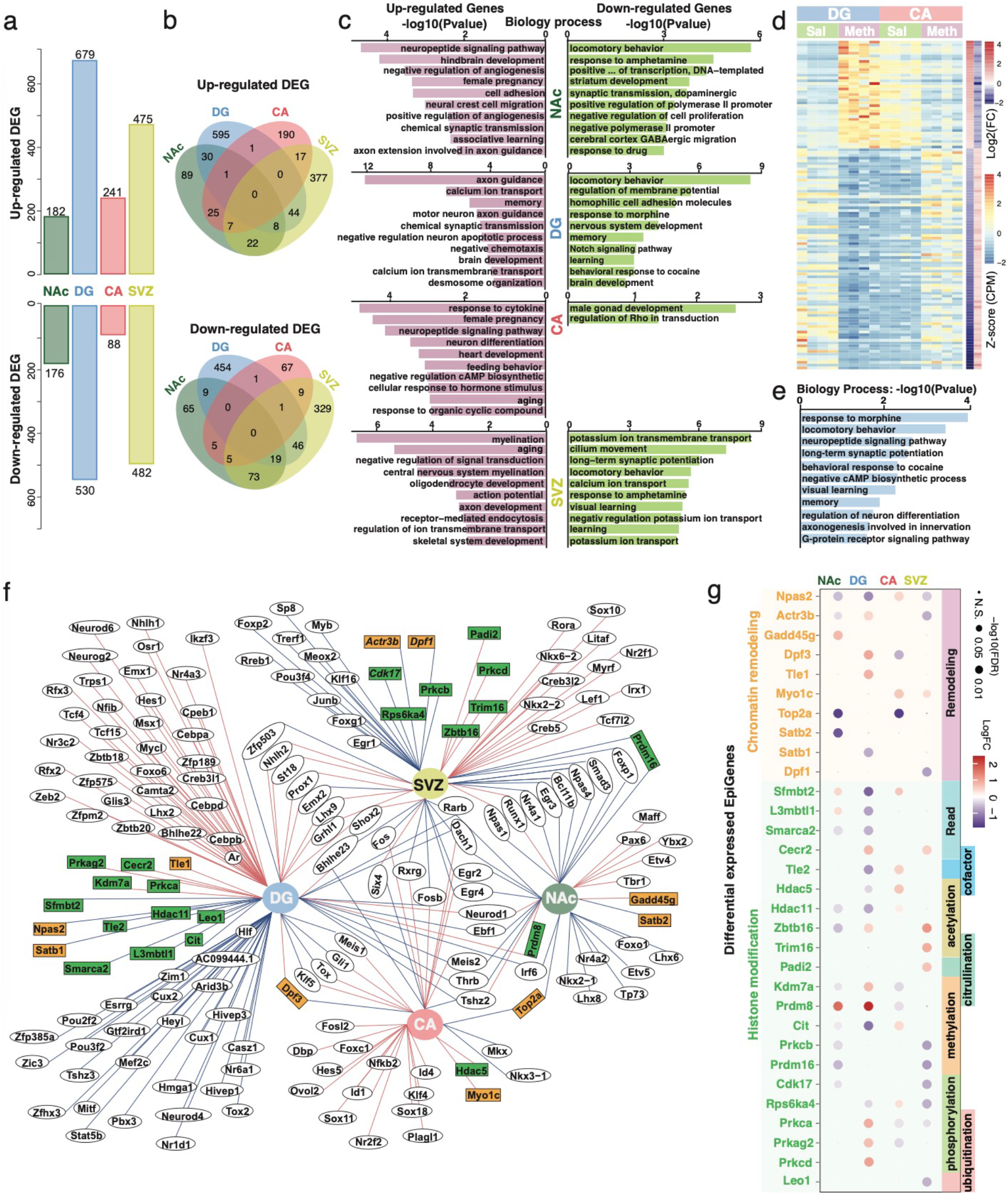
Differentially expressed genes in 4 rat brain regions in response to METH-overdose stimulus. **a**, Number of up- and downregulated differentially expressed genes (DEGs) in 4 rat brain regions. The largest number of DEGs was identified in the DG region. **b**, The number of shared up- and downregulated DEGs across 4 brain regions. Only a few DEGs were simultaneously affected by METH overdose in multiple regions. **c**, Gene Ontology (biological process) enrichment analysis of up- and downregulated DEGs in 4 brain regions. **d**, The DEGs showing opposing regulation patterns between the DG and CA. A total of 39 upregulated DEGs in the DG were identified as downregulated DEGs in the CA, and 80 downregulated DEGs in the DG were identified as upregulated DEGs in the CA. **e**, GO enrichment analysis of DEGs showing opposing regulation patterns between the DG and CA. These DEGs were highly enriched in neurological process terms. **f**, METH overdose induced significant expression changes in 146 transcription factors (TFs) and 31 epigenetic factors (epigenes) in 4 brain regions. The red line shows upregulated DEGs, and the blue line shows downregulated DEGs. Genes shown in circles are TF genes, and genes shown in squares are epigenes. The orange and green background colors match the chromatin remodeling and histone modification functions, respectively, in panel g. **g**, Differentially expressed epigenes with chromatin remodeling and histone modification functions in 4 brain regions.

The DG and CA are spatially connected regions in the hippocampal trisynaptic circuit and together contribute to new memory formation. However, we only found two commonly upregulated and two commonly downregulated DEGs after the METH-overdose stimulus in the DG and CA regions (Fig. 2b). Surprisingly, we found that 39 upregulated DEGs in the DG were identified as downregulated DEGs in the CA, and 80 downregulated DEGs in the DG were identified as upregulated DEGs in the CA (Fig. 2d and Additional file 2: Figure S2b). These types of DEGs were highly enriched in neurological processes, including response to morphine and cocaine, memory, visual learning, and long-term synaptic potentiation (Fig. 2e and Additional file 2: Figure S2c). Such opposite gene regulation in the DG and CA indicated a distinct regulation in the trisynaptic circuit in response to the METH-overdose stimulus.

Transcription factors (TFs) and epigenetic modification factors (epigenes) are critical components in the gene regulatory network. We precisely checked the TFs and epigenes associated with the METH-overdose stimulus in all four rat brain regions. In total, we found 146 TFs and 31 epigenes that significantly changed expression after the METH-overdose stimulus (Fig. 2f and Additional file 5). Twenty-seven TFs changed their expression in more than one brain region. Only six TFs, including *Egr2, Egr4, Rarb, Bhlhe23, Meis2*, and *Tshz2*, were differentially expressed in more than two brain regions, and *Dach1* was the only TF that responded to the METH-overdose stimulus in all four rat brain regions. Among 31 differentially expressed epigenes, 10 epigenes are considered to play essential roles in chromatin remodeling, and 21 genes are associated with histone modification, including acetylation, methylation, and phosphorylation (Fig. 2g). Three epigenes, including *Prdm8, Dpf3*, and *Top2a*, were found to be significantly changed in more than one brain region (Fig. 2g). For example, *Prdm8*, a conserved histone methyltransferase that acts predominantly as a negative regulator of transcription [54], was commonly upregulated in the NAc and DG, suggesting that epigenetic regulation might be involved in the repression of genes that respond to addictive substances in these two areas at 24 h after METH overdose (Fig. 2c and Additional file 2: Figure S2d). We further checked the expression changes of these epigenes across the four brain regions and found that many of these epigenes showed changes in expression. The *Hdac5* gene, which is responsible for the deacetylation of lysine residues on the N-terminal part of the core histones, showed an opposite regulation pattern between the DG and CA (Additional file 2: Figure S2e) [19]. The significant differential expression of epigenes in response to METH overdose could also result in epigenetic changes.

### METH overdose induced region-specific epigenetic changes in the rat brain

To further explore how METH-overdose stimulus remodels the epigenetic landscape in the rat brain, we carefully examined the alteration of chromatin accessibility in all four brain regions. In total, we identified 25,598 significant differential accessible regions (DARs) in the four rat brain regions responding to METH overdose; of these, 10,711 genomic loci became more accessible, and 17,816 regions became less accessible (Fig. 3a, Additional file 2: Figure S3a and Additional file 7). Similar to the case for the transcriptomic changes, the DG suffered the most significant changes in chromatin accessibility among all four regions. The 12,265 regions in the DG lost chromatin accessibility after the METH-overdose stimulus, suggesting that the DG region was the primary effect target of METH among the four regions we examined (Fig. 3a). Among DARs responding (becoming more or less accessible) to METH overdose, 89% of DARs were only identified in a single brain region, suggesting a solid region-specific epigenetic reaction to METH overdose in the four brain regions (Fig. 3b). The GO enrichment of genes around DARs indicated distinct molecular biological processes in the four brain regions (Additional file 2: Figure S3b and Additional file 8). In the NAc and DG, METH overdose induced lower accessibility in regions enriched in neurological functions, including synaptic plasticity, neurotransmitter transport, learning, and memory.

**Fig. 3.**
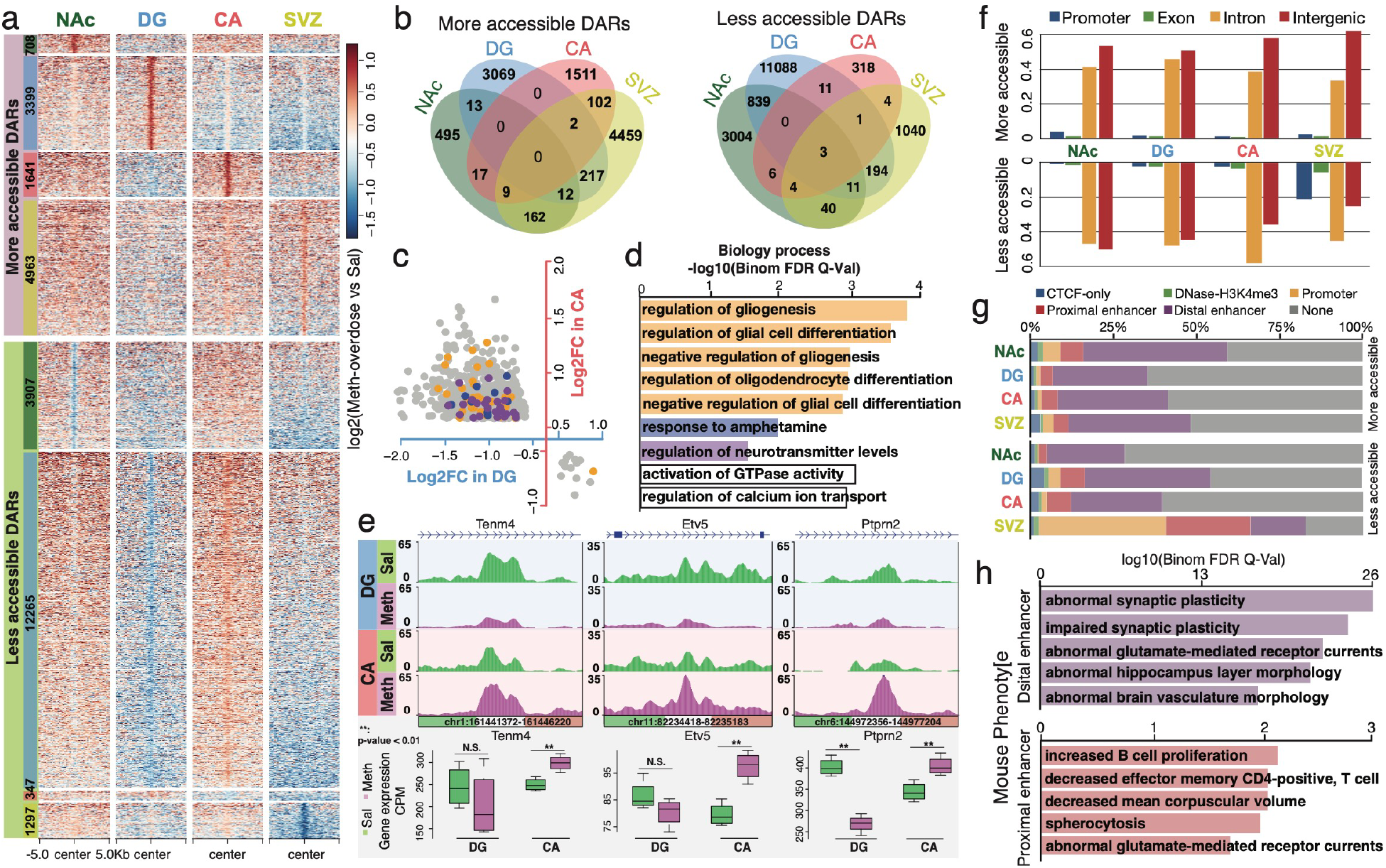
Differentially accessible regions in 4 brain regions induced by METH overdose. **a**, More and less accessible DARs identified in 4 brain regions in response to METH-overdose stimulus. Each dot represents a 500 bp window with log2 ratio of ATAC-seq signals by METH-overdose vs. Sal samples. **b**, The number of shared more and less accessible DARs across 4 rat brain regions. Eighty-nine percent of DARs were identified only in a single brain region. A small number of DARs were identified in multiple regions. **c**, The DARs showing opposite accessibility changes in the DG and CA in response to METH overdose. A total of 764 more accessible DARs in the CA were identified as less accessible DARs in the DG, and another 16 less accessible DARs in the CA showed more accessibility in the DG. Each dot represents one DAR. The DARs shown in orange, blue and purple color match the corresponding enriched biological process terms in panel d. **d**, GO enrichment analysis of DARs with opposite accessibility changes in the DG and CA. The genes around those DARs were significantly enriched in gliogenesis and glial cell differentiation (orange), response to amphetamine (blue) and regulation of neurotransmitter levels (purple). **e**, Three examples of DARs with opposite changes in accessibility in the DG and CA. Those DARs were located in the introns of three genes that had the same expression changes in the DG and CA. **f**, Genomic distribution of more and less accessible DARs in the rat genome (rn6). More than 90% of DARs were located in introns and intergenic regions of the rat genome. **g**, Distribution of rat–mouse ortholog DARs in cis-regulatory elements of the mouse genome (mm10), including CTCF-only, DNase-H3K4me3, promoter, proximal enhancer and distal enhancer. Approximately 30% of rat–mouse orthologous DARs showed enhancer function, including proximal enhancers and distal enhancers. Rat–mouse ortholog DARs: orthologous regions of rat brain DARs in the mouse genome (mm10). **h**, Enriched phenotypes of rat–mouse ortholog DARs annotated with distal enhancers and proximal enhancers in mouse.

Interestingly, the regulatory elements that gained chromatin accessibility in all four brain areas were highly enriched in nonneuronal functions. In the DG area, epigenetic changes under the METH-overdose stimulus were enriched in axon ensheathment, myelination, and oligodendrocyte and glial cell development. In the SVZ area, OCRs around immune-related genes were more open under the METH-overdose stimulus and enriched in myeloid leukocyte differentiation and endothelial cell chemotaxis (Additional file 2: Figure S3b). Such evidence suggests a potential hazard to glial cells after METH overdose.

Similar to the oppositely-regulated DEGs between the DG and CA (Fig. 2d), such opposite regulation was also observed at the epigenetic level after the METH-overdose stimulus. Compared to two more accessible DARs and 15 less accessible DARs that were commonly shared between the DG and CA, 47% of more accessible DARs in the CA (764) were identified as less accessible DARs in the DG responding to the METH-overdose stimulus, and another 16 of less accessible DARs in the CA showed more accessibility in the DG (Fig. 3c and Additional file 2: Figure S3c, S3d). The genes around those DARs with opposite changes in chromatin accessibility between the DG and CA were significantly enriched in gliogenesis and glial cell differentiation, response to amphetamine, and regulation of neurotransmitter levels (Fig. 3d and Additional file 2: Figure S3e). We identified an enhancer located in the intron of *Tenm4*, a gene that is associated with the establishment of proper connectivity within the nervous system [55], that exhibited significantly reduced chromatin accessibility in the DG after METH overdose (Fig. 3e) and increased chromatin accessibility in the CA area, accompanied by matched gene expression changes in these two subareas of the hippocampus (Fig. 3e). Similar epigenetic changes could also be observed in the intron regions of *Etv5* and *Ptprn2* (Fig. 3e), genes that could be essential for neuronal differentiation and required for normal accumulation of neurotransmitters in the brain [56, 57].

Approximately 90% of DARs associated with METH-overdose stimulus were located in intragenic or intergenic regions, except for the less accessible DARs in SVZ, suggesting that the distal regulatory elements, such as enhancers, were the primary targets that were affected by the METH-overdose stimulus (Fig. 3f and Additional file 9). To further explore the potential functions of DARs responding to METH overdose, we first identified the orthologous regions of DARs in the mouse genome. We then checked their regulatory potential using cis-regulatory element annotation in the mouse genome from ENCODE (CTCF-only, DNase-H3K4me3, promoter, proximal enhancer, and distal enhancer) [58]. On average, nearly 40% of the mouse orthologous regions of DARs responding to METH-overdose stimulus are explicitly annotated as regulators in the mouse genome, and ∼30% of them were annotated to have either proximal or distal enhancer function (Fig. 3g and Additional file 9). Interestingly, genes associated with two different enhancer DARs were enriched in distinct mouse phenotypes: genes associated with the distal enhancers were enriched in abnormal synaptic plasticity phenotypes, and genes associated with the proximal enhancers were mainly related to immunological phenotypes (Fig. 3h and Additional file 10).

To better understand the potential biological functions of DARs responding to METH overdose, we further explored the evolutionary conservation of DARs in the human and mouse genomes. Genome-wide alignment identified that the majority of the chromatin accessible regions in the rat brain were highly conserved and had orthologous counterparts in the mouse genome (90%) and the human genome (65%), in contrast to the relatively low conservation at the genome level (70% conservation between rat and mouse genomes, 30% conservation between rat and human genomes) (Fig. 4a, Additional file 2: Figure S4a-e and Additional file 11). We found that ∼70% of DARs associated with METH-overdose stimulus were conserved across all three species (rat-mouse-human DARs); 24% of DARs were rodent-specific (rat-mouse DARs), and less than 10% of DARs were rat-specific (rat-only DARs) (Fig. 4b). Furthermore, genes near (within 20 kb) the DARs with specific conservation statuses were enriched in specific biological process terms (Fig. 4c and Additional file 2: Figure S4b, S4c): genes associated with the most conserved type (rat-mouse-human DARs) were highly enriched in brain function and neurological process. The genes related to rodent-specific DARs were enriched in housekeeping functions, such as protein modification, DNA damage response, and protein nuclear transportation. However, the genes associated with rat-specific DARs were more enriched in immune-related biological processes (Fig. 4c).

**Fig. 4.**
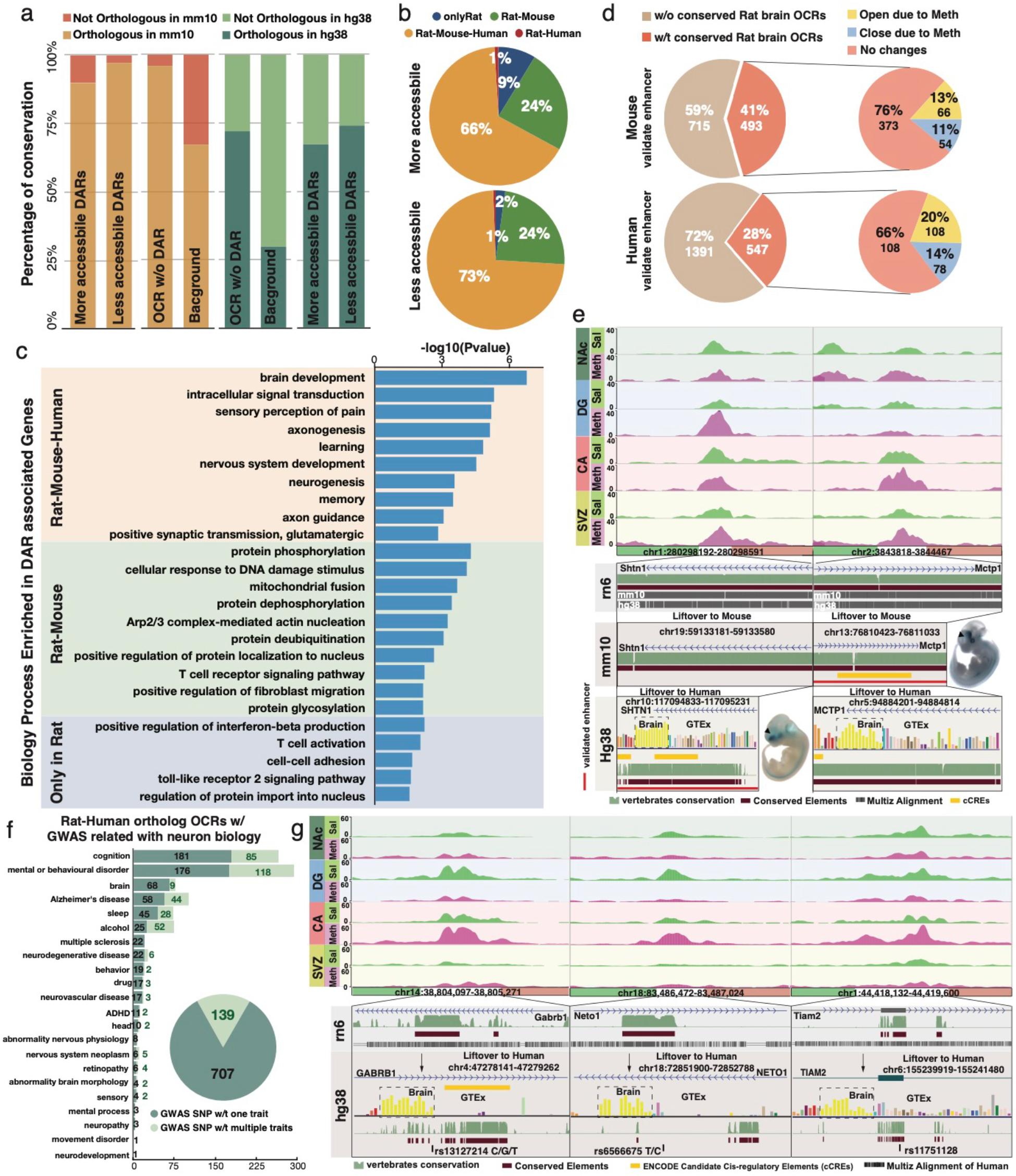
Evolutionary conservation of rat brain DARs with METH-overdose stimulus. **a**, Percentages of DARs, open chromatin regions (OCRs) and background regions with orthologous counterparts in mouse and human genomes (mm10 and hg38). **b**, Percentages of more and less accessible DARs with orthologs in the rat, mouse and human genomes. Approximately 70% of DARs were conserved across the three species (rat–mouse–human DARs); 24% of DARs were rodent-specific (rat–mouse DARs), and less than 10% of DARs were rat-specific (rat-only DARs). **c**, Biological process terms enriched in genes around (within 20 kb) DARs of three different conservation statuses: rat–mouse–human DARs, rat–mouse DARs and rat-only DARs. **d**, Percentages of mouse and human validated enhancers with ortholog sequences with OCRs in the rat brain. **e**, Two examples of DARs with orthologous interactions with human and mouse validated enhancers. **f**, Number of variants associated with neurologic phenotypes in genome-wide association studies (GWAS SNPs) that intersected with rat–human ortholog OCRs. **g**, Three examples of rat–human ortholog DARs containing GWAS SNPs associated with neuron biology.

Since most DARs associated with the METH-overdose stimulus were far from gene promoters (Fig. 3f), we further explored the potential enhancer functions of those DARs. Comparison with experimentally validated enhancers in mouse and human genomes [59] indicated that 493 mouse enhancers and 547 human enhancers had orthologous sequences with rat chromatin accessible regions (Fig. 4d and Additional file 12). Furthermore, 120 mouse enhancers and 186 human enhancers were conserved with DARs associated with METH-overdose stimulus (Fig. 4d). These experimentally validated enhancers targeted many essential genes in neuronal cells. For example, a rat-human ortholog DAR in the intron of the *Shtn1* gene (rn6, chr1:280298192-280298591) was more accessible in the DG after METH-overdose stimulus (Fig. 4e). In the human genome, this highly conserved ortholog region (hg38, chr10:117094833-117095231) is located in the center of one validated human enhancer (hg38, chr10:117094613-117095732), which was activated in the forebrain and midbrain of transgenic mice at embryonic Day 11.5 (Fig. 4e). The human *SHTN1* gene is highly expressed in brain regions and involved in generating internal asymmetric signals required for neuronal polarization and neurite outgrowth [60]. Another DAR (chr2:3843818-3844467) was located in the intron of *Mctp1*, an essential gene for the stabilization of neurotransmitter release and long-term maintenance of presynaptic homeostasis [61]; this site became more accessible in the CA and SVZ after METH-overdose stimulus (Fig. 4e). The mouse ortholog region of this DAR was located in the center of one validated enhancer (mm10: chr13:76809879-76811879) that was activated in the forebrain and hindbrain of transgenic mice at embryonic day 11.5 (Fig. 4e).

Previous studies showed that disease-associated genetic variants were enriched in regulatory elements [62-64]. We next studied the enrichment of phenotype-associated variants from genome-wide association studies (GWAS) of diverse traits and disorders collected by GWAS Catalog [65] in the conserved ortholog regulatory regions in the rat brain. In total, 846 SNPs associated with different neurological functions and disorders were located in 814 human–rat ortholog regions that were accessible chromatin regions in the rat brain (Fig. 4f, Additional file 2: Figure S4f and Additional file 13). A total of 707 SNPs were only associated with a single trait, and the remaining 139 SNPs were associated with multiple traits significantly enriched in cognition and mental/behavioral disorders. The SNP rs13127214 was associated with the unipolar depression field [60] and was mapped to *GABRB1*, a gene related to inhibitory synaptic transmission in the vertebrate brain [66]. SNP rs13127214, located in the human–rat conserved region (rn6, chr14:38,804,097-38,805,271), became close in the DG but open in the CA and SVZ after METH-overdose stimulus (Fig. 4g. left). We also found that the intronic SNP rs6566675 in *NETO1*, which plays critical roles in spatial learning and memory [67, 68], is located in the DAR chr18:83,486,472-83,487,024, which is closed in the NAc and DG but opened in the CA after METH stimulus (Fig. 4g. middle). The DAR chr1:44,418,132-44,419,600, closed in the NAc and DG but open in the CA, overlapped with the exon of the *Tiam2* gene. After alignment to the human genome, an orthologous region (chr6:155,239,919-155,241,480) was located within the *TIAM2* gene, which is highly expressed in the human brain and may play a role in the neural cell development [69, 70]. This conserved regulatory region contained a GWAS SNP (rs11751128) associated with the human intelligence [67].

### METH overdose induced a distinct gene regulatory network in the rat brain

To further explore the molecular regulation underlying the DAR responses to METH-overdose stimulus, we performed transcription factor (TF) binding motif enrichment analysis for both more and less accessible DARs in four rat brain regions. The binding motifs of many neuronal function-related TFs, including *Sox, Egr*, and *NeuronD1*, were highly enriched in the DARs associated with METH-overdose stimulus (Fig. 5a, Additional file 2: Figure S5a-c and Additional file 14, 15). We found 2434 (62.3%) less-accessible DARs in the NAc containing *Egr* binding sites, and genes around these *Egr* binding DARs were significantly enriched in neurological biology terms, such as memory, learning, and synaptic plasticity (Fig. 5b and Additional file 2: Figure S5b). We further built a protein-protein interaction (PPI) network for the genes in these enriched biological process terms in the NAc (Fig. 5b and Additional file 16). In the PPI network, *Rac1, Rac2, Drd1, Drd2, Camk2a, Camk2b, Gria1, Grin2a*, and *Fos* were found to be highly connected to other *Egr* targets, suggesting that *Egr* might regulate these network hubs in response to METH-overdose stimuli in the NAc region. *Rac1* is well known to be associated with cocaine addictive behavior [71, 72], and *Camk2a* is considered to be involved in the loss of control of ethanol consumption and cocaine dependence [73, 74]. *Gria1, Grin2a*, and *Fos* are highly implicated in drug addiction behavior and synaptic plasticity [75-78].

**Fig. 5.**
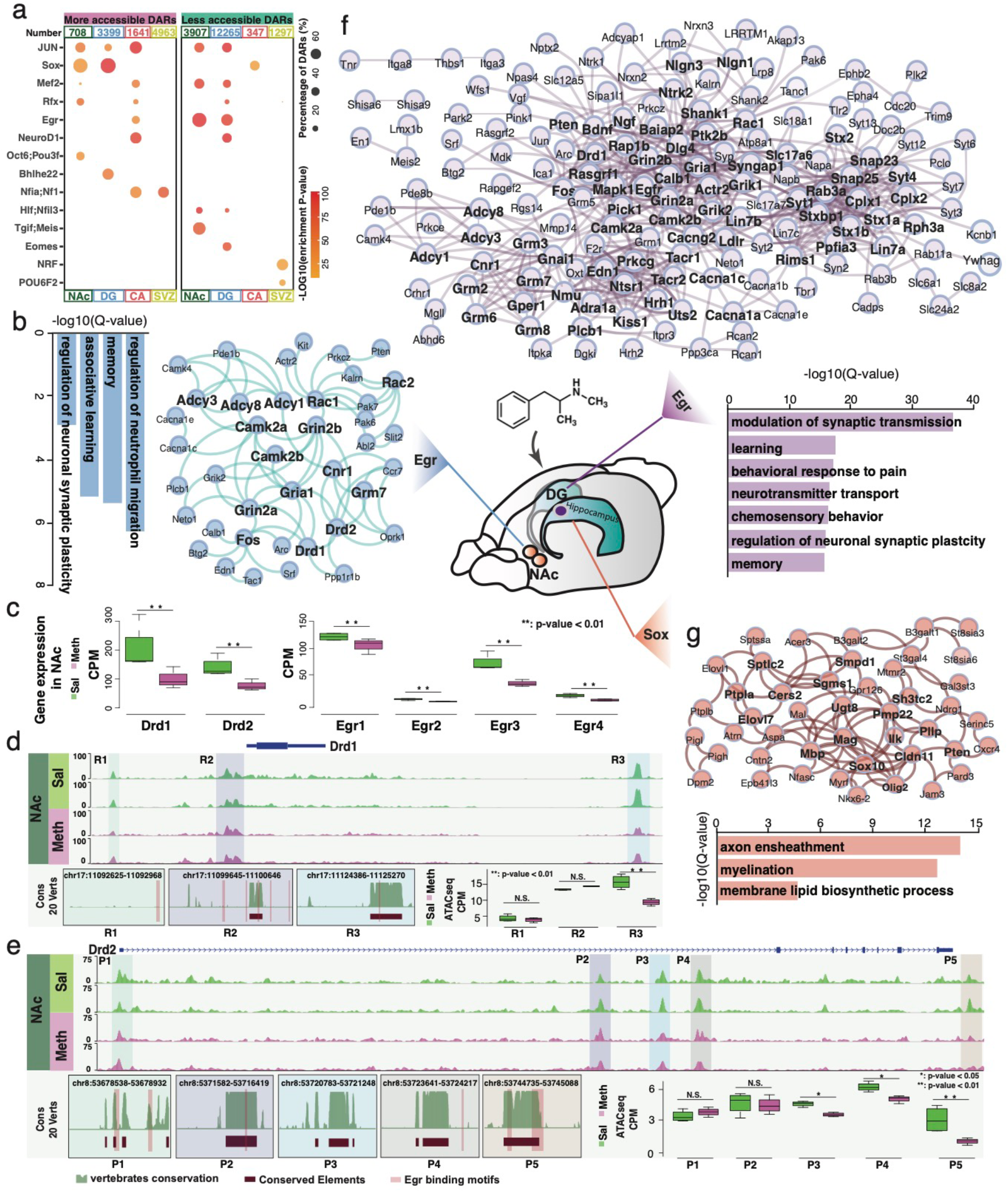
Distinct gene regulatory networks in 4 rat brain regions responding to METH overdose. **a**, Transcription factor (TF) binding motifs enriched in more and less accessible DARs of 4 rat brain regions in response to METH overdose. The size of the circle represents the percentage of DARs containing that TF binding motifs. The color scale represents the enrichment p-value. **b**, Enriched biological process terms of DARs with Egr binding motifs in the NAc and the gene regulatory network built by DAR-associated genes. **c**, Gene expression of *Drd1/2* and *Egr* family genes in the NAc in response to METH overdose. The OCRs around *Drd1* (**d**) and *Drd2* (**e**) in the NAc with the following analysis results: 20-vertebrate conservation of OCR, *Egr* binding motifs in OCR and open chromatin signals of OCR in METH-overdose and Sal samples. Enriched biological process terms and gene regulatory networks for DARs with *Egr* binding motifs (f) and DARs with *Sox* binding motifs (g) in the DG.

We specifically focused on the *Drd1* and *Drd2* genes, which are critical dopamine receptors in the NAc region and play central roles in responding to addictive behaviors for most drugs [79]. After the METH-overdose stimulus, we noticed that the downregulation of *Drd1* and *Drd2* was highly correlated with reduced expression of the *Egr* family (Fig. 5c and Additional file 2: Figure S5d). Three open chromatin regions around *Drd1*, including the promoter and potential enhancer regions, all contain *Egr* binding sites (Fig. 5d). We found that the 3’ downstream enhancer OCR (R3) lost open chromatin signals significantly after METH-overdose stimulus. The EGR binding motif in this METH-overdose-associated OCR was conserved between humans and rats (Fig. 5d). Similarly, three (P3, P4, and P5) of five OCRs around the *Drd2* gene significantly lost open chromatin signals after the METH-overdose stimulus. All five OCRs contained conserved elements between the human and rat genomes and harbored *Egr* binding motifs (Fig. 5e). Such results suggested that *Egr* might regulate *Drd1* and *Drd2* with a similar mechanism in both rat and human brains.

In the DG region, 5088 DARs were found to contain the *Egr* binding motif, which accounted for 41.5% of the total open chromatin regions that lost open chromatin signals after the METH-overdose stimulus (Fig. 5a, Additional file 2: Figure S5a and Additional file 15). Genes around these less opened DARs were highly enriched in synaptic transmission, learning, and behavioral response to pain (Fig. 5f, Additional file 2: Figure S5b and Additional file 16). In the PPI network of these *Egr* targets in the DG region, the top key nodes tightly associated with distinct psychiatric disorders and addictive behaviors in previous studies included *Syt1, Stx1a, Dlg4, Cplx1, Gria1, Snap25, Rab3a*, and *Bdnf* [75, 80-85]. Our results suggested that these essential genes could all be regulated by the *Egr* family and emphasize the importance of *Egr* in the rat DG region in response to METH-overdose stimulus.

Meanwhile, we found that the Sox binding motif was highly enriched in the METH-overdose-induced more-open DARs in both the NAc and DG regions (Fig. 5a). A total of 62.4% of NAc more-open DARs (442) and 70% of DG more-open DARs (2,380) contained *Sox* binding motifs (Additional file 2: Figure S5a and Additional file 15). The genes around these *Sox* binding DARs in the DG region were highly enriched in axon ensheathment, myelination, and membrane lipid biosynthetic processes (Fig. 5g, Additional file 2: Figure S5b and Additional file 16). PPI network analysis suggested that *Mbp, Sox10, Mag, Cldn11, Ugt8, Pmp22*, and *Cers2* were the top connected nodes with critical regulatory roles in myelination and lipid metabolism and ensheathment in the oligodendrocytes [86-89].

## Discussion

Methamphetamine (METH) is a stimulant amphetamine drug that is extremely persistently addictive, with 61% of individuals treated for METH use disorder relapsing within one year [16-18]. Recently, developed animal models, especially rodent models, helped us better understand the molecular consequences of substance abuse [1, 23, 27]. Although previous studies illustrated that METH exposure could cause dramatic epigenetic changes in the NAc and frontal cortex [1, 3, 27, 33, 44], there is still a lack of a clear description of the molecular changes in different brain regions under METH exposure.

Here, in addition to the most frequently studied region, the NAc, we explored the METH-induced epigenetic and transcriptomic changes in other areas of rat brains, namely the dentate gyrus, Ammon’s Horn, and subventricular zone. This work represents the most comprehensive dataset to date of METH-induced transcriptome and chromatin accessibility from multiple rat brain regions. Surprisingly, METH exposure induced genome-wide epigenetic alterations and resulted in dynamic gene expression changes in all four brain regions. More importantly, the molecular changes induced by METH were rarely shared among different areas, suggesting that different brain regions respond to METH exposure in a highly region-specific fashion. For example, approximately one thousand genes underwent expression changes in response to METH exposure in the DG and SVZ regions, while only approximately three hundred genes underwent expression changes in the NAc and CA regions (Fig. 2a). This large difference in transcriptional response does not correspond with the sizes of the sets of genes with region-specific expression overall, which number only in the hundreds for each region (Fig. 1d). All these results suggest that the same genes have distinct regulatory mechanisms in different brain regions. We also noticed substantial expression alterations in many genes associated with histone modification and chromatin remodeling after METH exposure in all four brain regions (Fig. 2f, g). This finding provides evidence that complicated epigenetic remodeling events at different levels can be induced by METH exposure, as previous studies have reported [1, 3, 19, 24, 27, 33, 45].

More specifically, we noticed that two subregions of the hippocampus, CA and DG, showed the opposite responses to METH exposure at both the transcriptomic (Fig. 2d) and epigenetic (Fig. 3c) levels. These oppositely regulated genes were highly associated with responses to different substances in the brain, memory, neuron, and glial cell differentiation (Fig. 2e, Fig. 3d). The opposite response between the CA and DG might be associated with the specific functionality of the two subregions. As a part of the hippocampal formation in the temporal lobe of the brain, the dentate gyrus is generally believed to contribute to the formation of new synapse connections and episodic memories [90-93]. The CA region of the hippocampus, also called Ammon’s horn, plays an important role in long-term memory [93, 94]. Thus, even short-term METH stimulation might create new synaptic connections in the DG region, affecting the long-term memory in the CA region. Moreover, in the DG region, we noticed that the open chromatin regions (OCRs) around genes associated with glial cell differentiation were less open after METH exposure (Fig. 3d), and the same OCRs became more open in the CA. This result suggested that the glial cells in the hippocampus might be more active and vulnerable to METH exposure.

We also performed a comparative genome analysis to understand the potential function of DARs identified in different brain regions. Over 70% of DARs associated with METH exposure were conserved between humans and rats. The genes around these DARs were highly enriched in fundamental functions of the brain, such as brain development, neurogenesis, learning, and memory (Fig. 4a, c). In contrast, genes around rodent- and rat-specific DARs were more often associated with protein modification and immune response; such results emphasize the unique species-specific characteristics of the rat model in addiction research. We also noticed many known SNPs associated with neuronal biology and disease located in the conserved human ortholog counterparts of DARs responding to METH exposure in the rat brain (Fig. 4f). Most importantly, we found that more than 300 of these DARs validated brain-specific activation (Fig. 4d, e). Thus, we firmly believe that the conserved human orthologs of DARs identified in this study are highly likely to play essential roles in human METH addictive behaviors.

We also constructed regional-specific regulatory networks in response to METH exposure in the rat brain (Fig. 5b, f, g). By using a genomic DNA context-based approach, we were able to connect crucial upstream TF regulators to their downstream target genes and eventually better understand the molecular response to METH exposure at an upstream regulatory level. For example, both *Drd1* and *Drd2* are critical players in the reward pathway directly associated with addiction to multiple substances, including METH [79]. Meanwhile, the *Egr* family is broadly affected by METH exposure [45, 95-97]. Our results provide new information regarding the changes in *Drd1/2* expression in response to METH exposure in the rat NAc region through the *Egr* family. Meanwhile, certain evidence indicates that METH exposure greatly affects the biological functions of glial cells [98-100]. In both the NAc and DG regions, our results indicated that *Sox* binding sites were highly enriched in the DARs associated with genes that regulate myelination and ensheathment, suggesting that *Sox* family members could be upstream regulators in glial cells responding to METH exposure.

## Conclusions

Our study investigated the transcriptomic and epigenetic reaction to METH overdose exposure of four rat brain regions, including the NAc, DG, CA, and SVZ. We discovered that METH overdose induced 2,254 differentially expressed genes (DEGs) and 25,598 differentially accessible regions (DARs) in the four rat brain regions in a solid region-specific response pattern at the transcriptomic and epigenetic levels. An interesting opposite regulation pattern to react METH overdose exposure was observed between the rat CA and DG regions for the first time. 70% of METH overdose-induced epigenetic perturbations in rat brains were orthologous conserved in the human genome at the DNA sequence level. In general, our study provided a valuable multi-omics resource to better understand the molecular changes of the brain after METH overdose exposure.

## Methods

### Animals

Adult male Sprague-Dawley rats (Harlan, Indianapolis, IN, USA) (weighing 250–300 g on arrival) were pair-housed under a 12 h light/dark cycle in a temperature-controlled (20–22 °C) and humidity-controlled room. Food and water were available ad libitum. The animals were allowed to acclimatize for one week before the start of the study. All animal procedures were conducted between 7:00 A.M. and 7:00 P.M. in strict accordance with the National Institutes of Health (NIH) Guide for the Care and Use of Laboratory Animals and were approved by the Institutional Animal Care and Use Committee (IACUC) at Wayne State University. The description of animal procedures meets the ARRIVE recommended guidelines described by The National Centre for the Replacement, Refinement and Reduction of Animals in Research [101].

### Methamphetamine administration

(+)-Methamphetamine hydrochloride (METH, 10 mg/kg free base) (Sigma-Aldrich, St. Louis, MO) or saline (1 mL/kg) was administered to the rats every 2 h in four successive intraperitoneal (i.p.) injections, as previous studies [102-104]. To measure hyperthermia, the core body temperatures of the rats were measured with a rectal probe digital thermometer (Thermalert TH-8; Physitemp Instruments, Clifton, NJ) before the beginning of the treatment (baseline temperatures) and at 1 h after each METH or saline injection. Rats were sacrificed by decapitation at 24 h after the last injection of the drug or saline.

### Library construction

Total RNA was isolated via TRIzol Reagent (Thermo Fisher Scientific, 15596026), Phasemaker Tubes (Thermo Fisher Scientific, A33248) and RNA Clean & Concentrator-5 (Zymo Research, R1013). In brief, rat brain tissues were homogenized in 1ml of TRIzol Reagent, the tissue lysates were transferred to a pre-spined Phasemaker Tube. 0.2ml of chloroform was added to the tube, the tubes were then shook for 15 seconds. The mixture was centrifuged at 16,000g for 5 minutes, the top aqueous phase was transferred to a microcentrifuge tube. An equal volume of ethanol was added to the aqueous phase. The RNA purification with DNase treatment was performed following the manual of the RNA Clean & Concentrator kit. Then the Ribosomal RNAs were removed from 500ng for the total RNA using the NEBNext rRNA Depletion kit (NEB, E6310). Skipping the mRNA isolation part, RNA-seq libraries were then constructed using 10ng of rRNA depleted total RNA with Universal Plus mRNA-seq kit (TECAN, 0520-A01) following the kit manual. 2×75bp paired-end sequencing was run for all libraries on the Illumina NextSeq 500 platform.

ATAC-seq was generated using the omni ATAC protocol for frozen tissues (Nature Methods volume 14, pages959–962, 2017). In brief, rat brain tissues were homogenized in 2ml of cold 1× homogenization buffer. Nuclei were layered from the tissue lysate with iodixanol solution. 50,000 nuclei were used in the transposition reaction with 100nM of Transposase (Illumina, 20034197). The ATAC-seq libraries were prepared by amplifying for 9 cycles on a PCR machine with NEBNext High-Fidelity 2× PCR master mix (NEB, M0541). 2×75bp paired-end sequencing was run for all libraries on the Illumina NextSeq 500 platform.

### Raw sequence data and processing

Total 32 RNA-seq fastq files and 29 ATAC-seq fastq files were generated from 4 rat brain regions, including Nucleus accumbens (NAc), Dentate gyrus (DG), Ammon’s horn (CA), and Subventricular zone (SVZ). For both RNA-seq and ATAC-seq, each region had 4 samples with saline treatment (Sal) and 4 samples with methamphetamine overdose (METH-overdose), except SVZ region only had 3 Sal samples and 2 METH-overdose samples for ATAC-seq data.

ATAC-seq data of 4 brain regions were separately processed by AIAP package that contained an optimized ATAC-seq data QC and analysis pipeline with default parameters[105]. Open chromatin regions (OCR) generated by AIAP were used in downstream analysis. Then, mergeBed was used to generate consensus OCRs of two conditions of 4 brain regions [106]. RNA-seq data were processed as in previous studies [107]. RNA-seq data of 4 rat brain regions were processed by Cutadapt (v2.7; --quality-cutoff=15,10 --minimum-length=36), FastQC (v0.11.4), and STAR (v2.5.2b; --quantMode TranscriptomeSAM --outWigType bedGraph -- outWigNorm RPM) to do the trimming, QC report and rat genome mapping (rn6) [108-110]. Then, gene expressions in rat brain regions was calculated by featureCounts (-p -T 4 -Q 10) based on UCSC gene annotation of rat [111, 112].

### Region-specific genes and OCRs in 4 regions of normal rat brain

EdgeR was used to identify region-specific genes that significantly highly expressed in one normal region comparing to other three regions (log2(fold change)>1 and FDR<0.01) [113, 114]. The GO enrichment analysis of region-specific genes in 4 regions were performed by DAVID (Database for Annotation, Visualization and Integrated Discovery, v6.8) [115, 116]. The region-specific OCRs were also identified in 4 rat brain regions with EdgeR method (log2(fold change)>1 and FDR<0.01) [113]. Then, the average signals of normal ATAC-seq samples in each region were generated by bigWigMerge and visualized by plotHeatmap of deepTools software [117]. The mouse ortholog regions of region-specific OCRs were generated by liftOver with parameter “-minMatch=0.6” and then those ortholog regions were used to identify enriched biological process terms in 4 brain region with GREAT (version 4.0.0) [112, 118]. The analysis settings of GREAT included that: 1)

Species assembly: Mouse, NCBI build 38; 2) Background regions: whole-genome; 3) Association rule: Basal plus extension. Then, top 20 enriched terms of biological process in 4 rat brain regions were filtered with cutoffs of Binom FDR Q-Val < 0.05 and Hyper FDR Q-Val < 0.05 simultaneously.

### METH-overdose induced differential expression genes in 4 rat brain regions

The significantly differential expression genes (DEGs) induced by METH-overdose in 4 rat brain regions vs saline samples were identified by EdgeR method with the cutoffs of log2FoldChange > 1 and FDR< 0.01. The Venn plot of intersection for up- and down-regulated DEG among 4 regions were respectively generated by jvenn [119]. The enriched terms of biology process were identified by DAVID separately for up- and down-regulated DEG. Next, the gene lists of transcription factors (TFs) and epigenetic factors (epigenes) were separately gathered from the AnimalTFDB3.0 and Epifactors databases [120, 121].

### Differential accessible regions identified in 4 rat brain regions due to METH-overdose

The EdgeR was used to identify significantly differential accessible regions (DARs) induced by METH-overdose in 4 brain regions comparing to saline samples with cutoffs of log2FoldChange > log2(1.5) and FDR< 0.001 [113]. The log2 values of the differential accessibility in 4 brain regions after METH-overdose stimulus were generated based on bigwig files by bigwigCompare and visualized by plotHeatmap of deepTools [117]. The Venn plots of intersection for both more and less accessible DARs among 4 brain regions were respectively generated by jvenn [119]. The liftOver was used to identify ortholog regions of rat brain DARs in mouse genome (mm10) with parameter “-minMatch=0.6” [112]. Then, those ortholog regions were used to iednitfy the top 20 enriched terms of biology process in 4 rat brain regions by GREAT with default parameter and cutoffs of Binom FDR Q-Val < 0.05 and Hyper FDR Q-Val < 0.05 [118]. Three examples of DARs with reversed accessibility between DG and CA were visualized by WashU Epigenome Browser. The intersectBed method was used to determine numbers of DARs across different genomic features (promoters, exons, introns, and intergenic regions), which were defined by using UCSC gene annotation of rat genome. The mouse ortholog regions of rat brain DARs intersected with mouse cis-regulation elements (CREss) from ENCODE database were used to explore potential regulatory function of those DARs [58]. Then, GREAT was used to identify enriched mouse phenotypes of those ortholog DARs with different CRE annotation types.

### Evolutionary conservation of DARs

The orthologous regions in mouse (mm10) and human (hg38) genomes for OCRs in rat brain regions were separately identified by using liftOver software with parameter “-minMatch=0.6” [65]. The rat genome was divided into 500bp windows as background regions and the orthologous conservation of background regions were measured as the same standard as OCRs described above. The rat-brain DARs were classified into 4 groups based on the ortholog in mouse and human genomes: Rat-Mouse-Human DARs were conserved across rat, mouse and human genomes; Rat-Mouse or Rat-Human DARs were DARs of rat had orthologous regions only in mouse or human genome; onlyRat DARs were not orthologous in mouse and human genomes. Then, those DARs form different groups were mapped to nearest genes within 20kb away, and the specific DAR-mapped genes of those 4 groups were separately used to identify the enriched terms of biology process by DAVID.

The experimentally validated human (hg19) and mouse (mm9) enhancers were downloaded from VISTA Enhancer Browser database and were separately transferred to the coordinates in hg38 and mm10 genomes by liftOver with parameter “”-minMatch=0.95” [59, 112]. Then, intersectBed method was used to separately identify the mouse and human validated enhancers that can be overlapped by mouse and human orthologous regions of rat brain OCRs. The two examples of DARs that had orthologous intersection with human and mouse validated enhancers were visualized by WashU Epigenome Browser [122]. The results of vertebrates-conservation, multiz-alignments, CREs and gene expression in GTEx data portal were generated by USCS genome browser [112]. The activation patterns of those two validated enhancers at E11.5 stage were also downloaded from VISTA Enhancer Browser.

The variants-trait information of genome-wide association studies (GWAS) were downloaded from GWAS Catalog (<https://www.ebi.ac.uk/gwas/home) >[65]. The GWAS SNPs associated with neurological process were identified based on mapped traits. Then, intersectBed was used to overlap those GWAS SNPs and rat-human orthologous OCRs. Three examples of rat-human orthologous DARs contained GWAS SNPs were visualized by WashU Epigenome Browser. Those three OCRs associated vertebrates-conservation, alignment, CREs and GTEx expression were generated from USCS genome browser.

### Motif enrichment of DAR and Gene regulation network

The transcription factors binding motifs (TFBS) enriched in more and less accessible DARs of 4 rat brain regions were separately analyzed by using findMotifsGenome.pl (-size given) of HOMER software [123]. The significantly enriched de novo binding motifs in 4 regions were identified with three conditions: 1) at least 5% of accessible DARs in one region contained the TFBS; 2) match score of TFBS should be greater than 0.85; 3) P-value of TFBS should be less than 1e-11. Then, known transcription factor (TF genes) under the enriched TFBS were extracted with match score > 0.85.

The DARs contained the Egr, Sox and NeuroD1 binding motifs were extracted from the HOMER results and ortholog regions in mouse of those DARs were used to identify enriched biological process terms with GREAT. The DARs associated genes in neurological biology process were extracted and used to build the gene regulation networks by String [124].

The open chromatin signals of OCRs around Drd1 and Drd2 genes in NAc region were visualized by WashU Epigenome Browser. The vertebrates-conservation of OCRs associated with Drd1 and Drd2 genes were generated by USCS genome browser. The FIMO software was used to scan TFBS motifs in those OCRs associated with Drd1 and Drd2 genes based on motif weigh matrix file (JASPAR_CORE_2016_vertebrates.meme) from JASPAR [125, 126].

## Authors’ Contributions

BZ, PM, and AM designed and supervised the study. VB and AM performed the animal experiment. XX generated transcriptomic and epigenomic sequencing data. BM processed data and performed analysis. BM, PM, AM, and BZ wrote the manuscript with input from all the other authors. All authors read and approved the final manuscript.

## Acknowledgements

We would like to thank Akhil Sharma for the technical assistance of the animal experiment.

## Consent for publication

Not applicable.

## Ethics approval and consent to participate

The animal experiments were followed with the National Institutes of Health (NIH) Guide for the Care and Use of Laboratory Animals and were approved by the Institutional Animal Care and Use Committee (IACUC) at Wayne State University. The description of animal procedures meets the ARRIVE recommended guidelines described by The National Centre for the Replacement, Refinement and Reduction of Animals in Research [101].

