## Additional file 2 for "Methamphetamine induced regional-specific transcriptomic and epigenetic changes in the rat brain"

**Supplementary Figures**

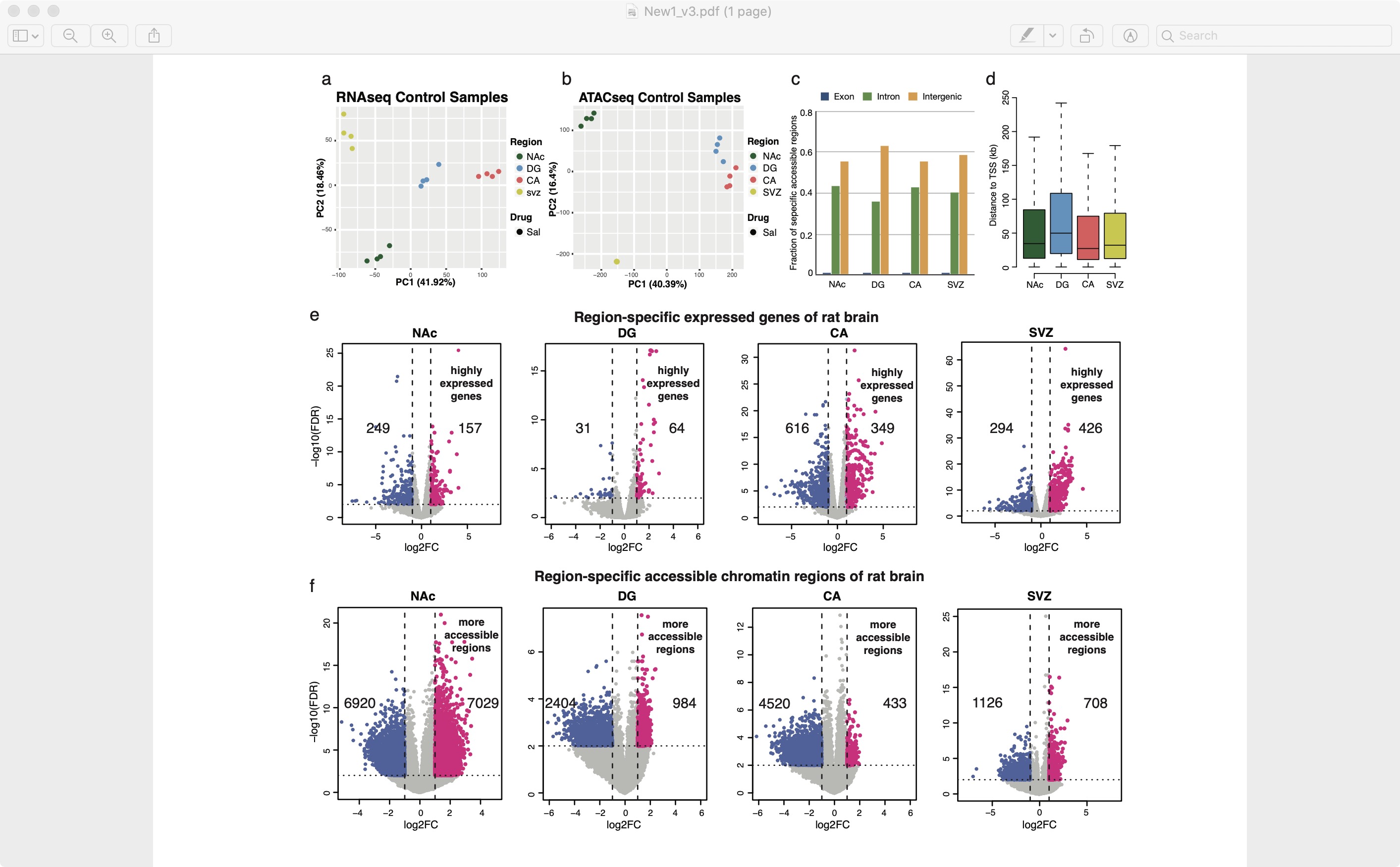

**Figure S1. RNA-seq and ATAC-seq data of 4 brain regions from control and METH-overdose rat samples.** The principal analysis (PCA) of RNA-seq data (a) and ATAC-seq data (b) from 4 normal rat brain regions. c, Genomic distribution of 4 rat brain region-specific accessible regions in rat genome: exon, intron, and intergenic regions. Large fraction of those specific accessible regions located in nitrogenic and intergenic regions. d, Boxplot about the distance to TSS of region-specific accessible regions in 4 rat brain regions. e-f, The volcano plots of region-specific expressed genes (e) and accessible chromatin regions (f) in 4 rat brain regions. Red dots represented the specific genes and accessible regions in each brain region. Log2FC: log2 value of fold changes with one normal region against other regions; -log10(FDR): -log10 value of false discover rate (FDR).

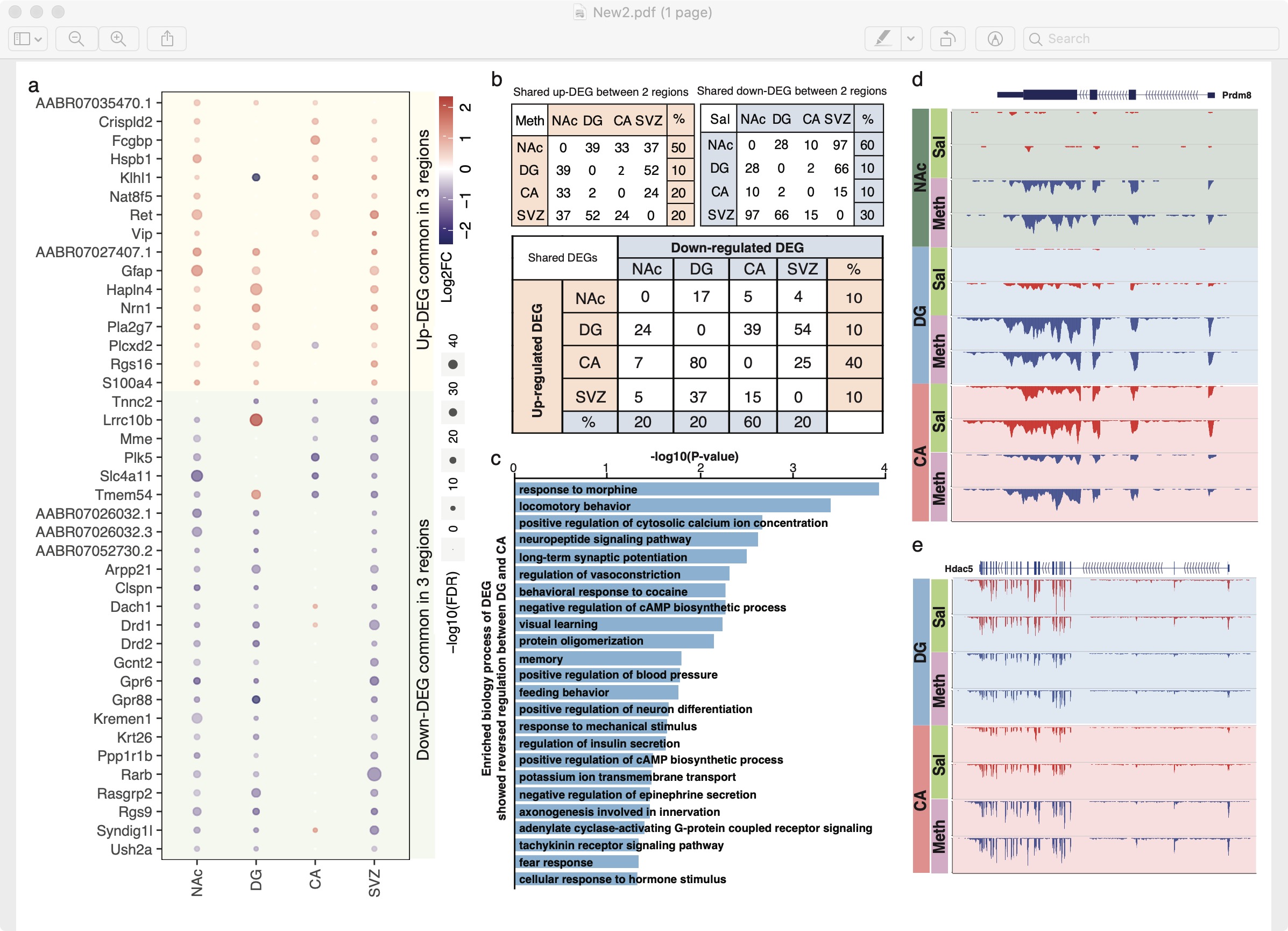

**Figure S2. The differential expressed genes (DEGs) identified in 4 rat brain regions responding to METH-overdose.** a, Up and down-regulated DEGs were common in at least 3 brain regions. b, Number of common genes between 2 regions separately for up and down-regulated DEGs, and the number of common DEGs between DG and CA showed reversed regulation pattern. Large percent of up and down-regulated DEGs in NAc were also identified in another one region. And large percent of DEG in CA regions showed reversed regulation pattern in another one region, especially in DG region. c, Biology processes of GO term enriched in the DEGs that showed reversed regulation between DG and CA regions. d-e, Examples of Prdm8 and Hdac5 genes showing differential expression in brain regions visualized by WashU Epigenome Browser.

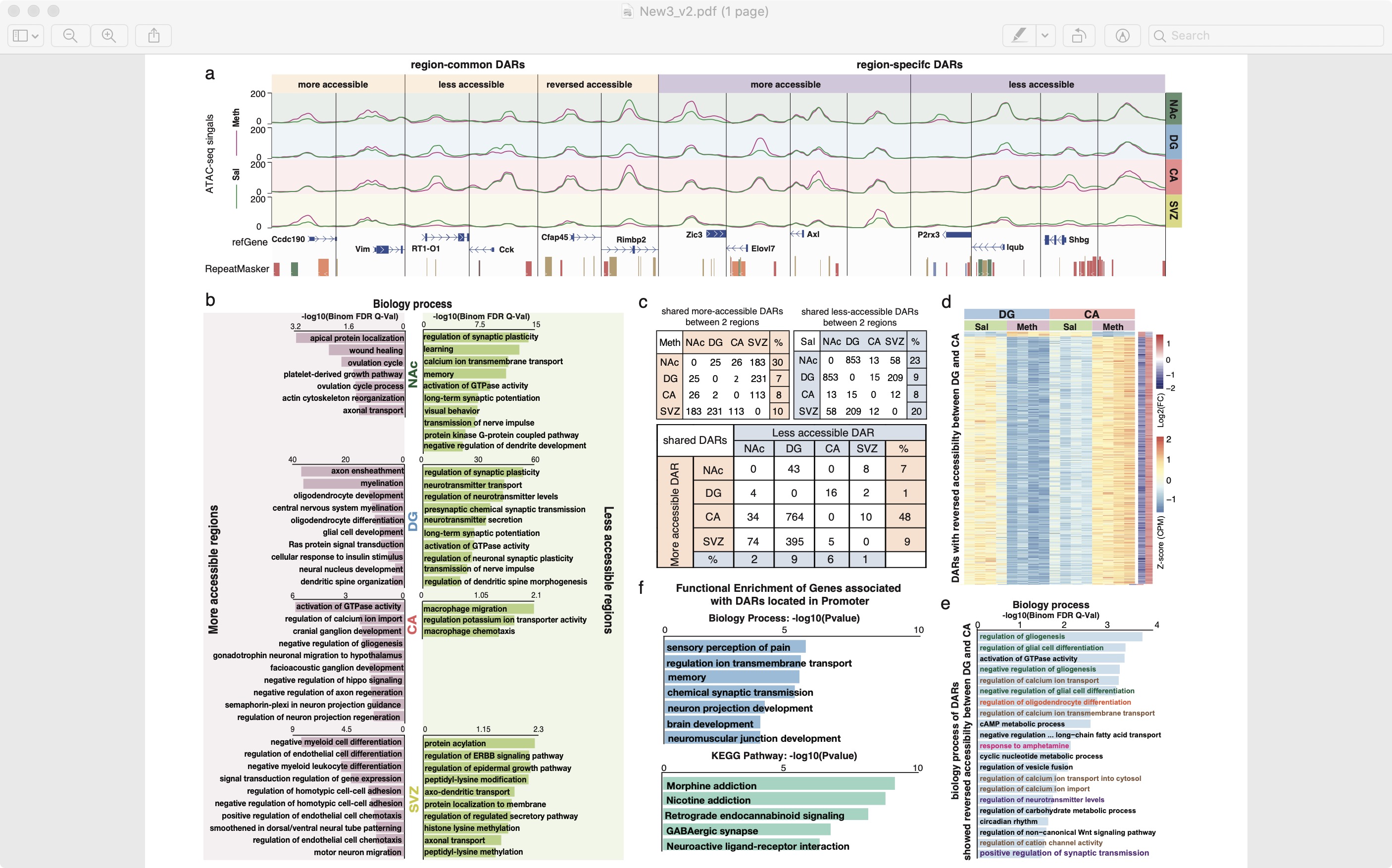

**Figure S3. The differential accessible regions (DARs) in rat brain regions after METH-overdose.** a, Examples of DARs showed common and specific accessibility in 4 rat brain regions. b, Enriched biology process of more and less accessible DARs in 4 rat brain regions. c, Number of common DARs between 2 regions separately for more and less accessible DARs, and the number of common DARs between DG and CA showed reversed accessibility. Large percent of more accessible DARs in CA region showed reversed accessibility in DG region. d, Heatmap about Z-score of DARs that displayed reversed accessibility between DG and CA. e, Biology processes significantly enriched in DARs showing reversed accessibility between DG and CA. f, Enriched biology processes and KEGG pathways based on associated genes of DARs located in the promoter regions.

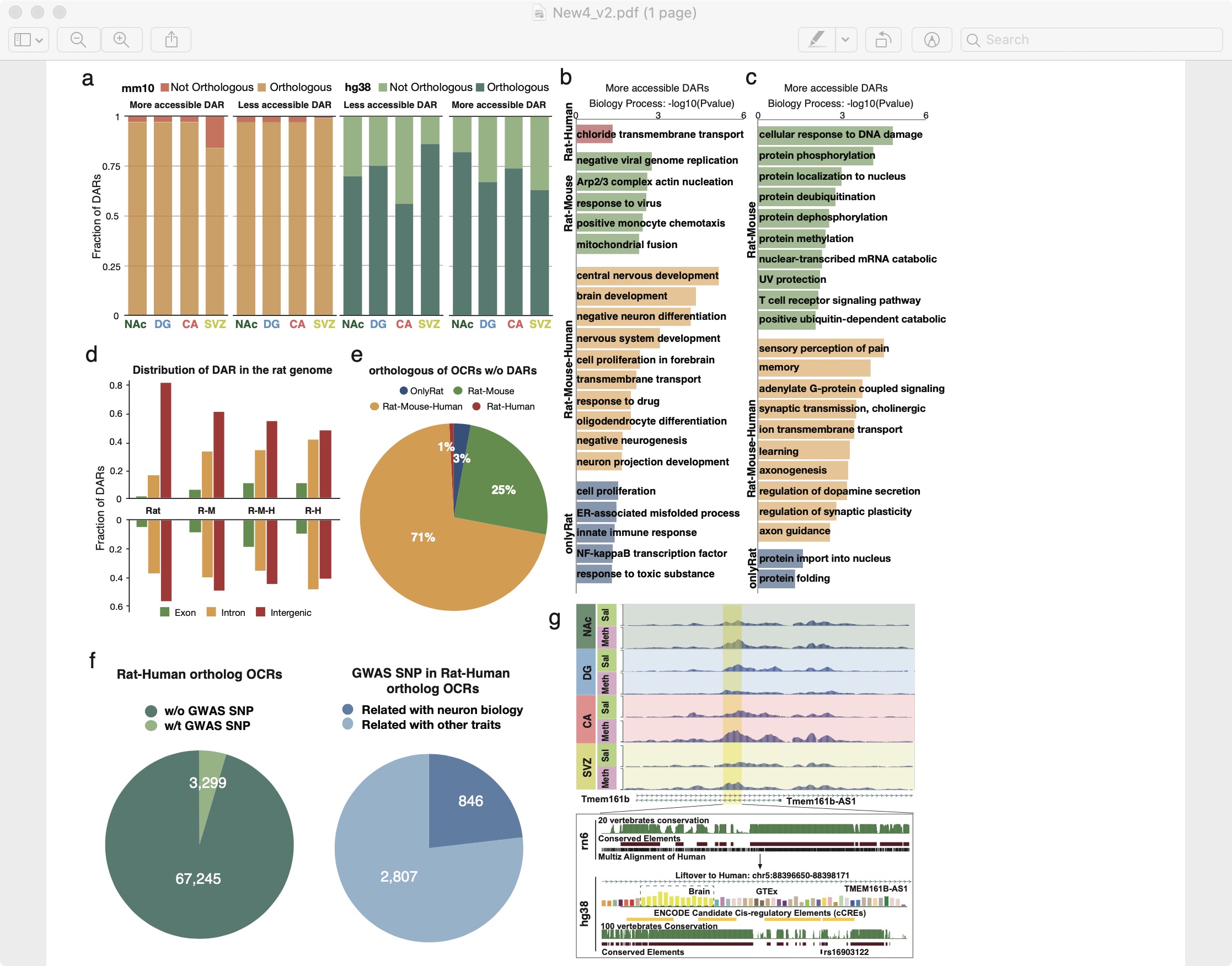

**Figure S4. Evolutionary conservation of Rat brain DARs responding to METH-overdose.** a, Percent of more and less accessible DARs had orthologous counterparts in mouse and human genome (mm10 and hg38). b-c, Biology processes enriched in genes around (within 20kb) DARs separately for more and less accessible DARs with different conservation status: rat-mouse-human (R-M-H), rat-human (R-H), rat-mouse (R-M) and rat-only (Rat). d, Genomic distribution of DARs from 4 different conservation status: exon, intron, and intergenic regions. Large fraction of DARs located in intergenic regions for all 4 groups. e, Percent of open chromatin regions (OCRs) without DARs from different conservation status. More than 70% of those OCRs were ortholog in rat, mouse, and human genome. f, Number of Rat-Human ortholog OCRs with variants from genome-wide association studies (GWAS SNP) and number of those GWAS SNP associated with neuron biology. g, Example of rat-human ortholog DAR overlap with GWAS SNPs associated with neuron biology.

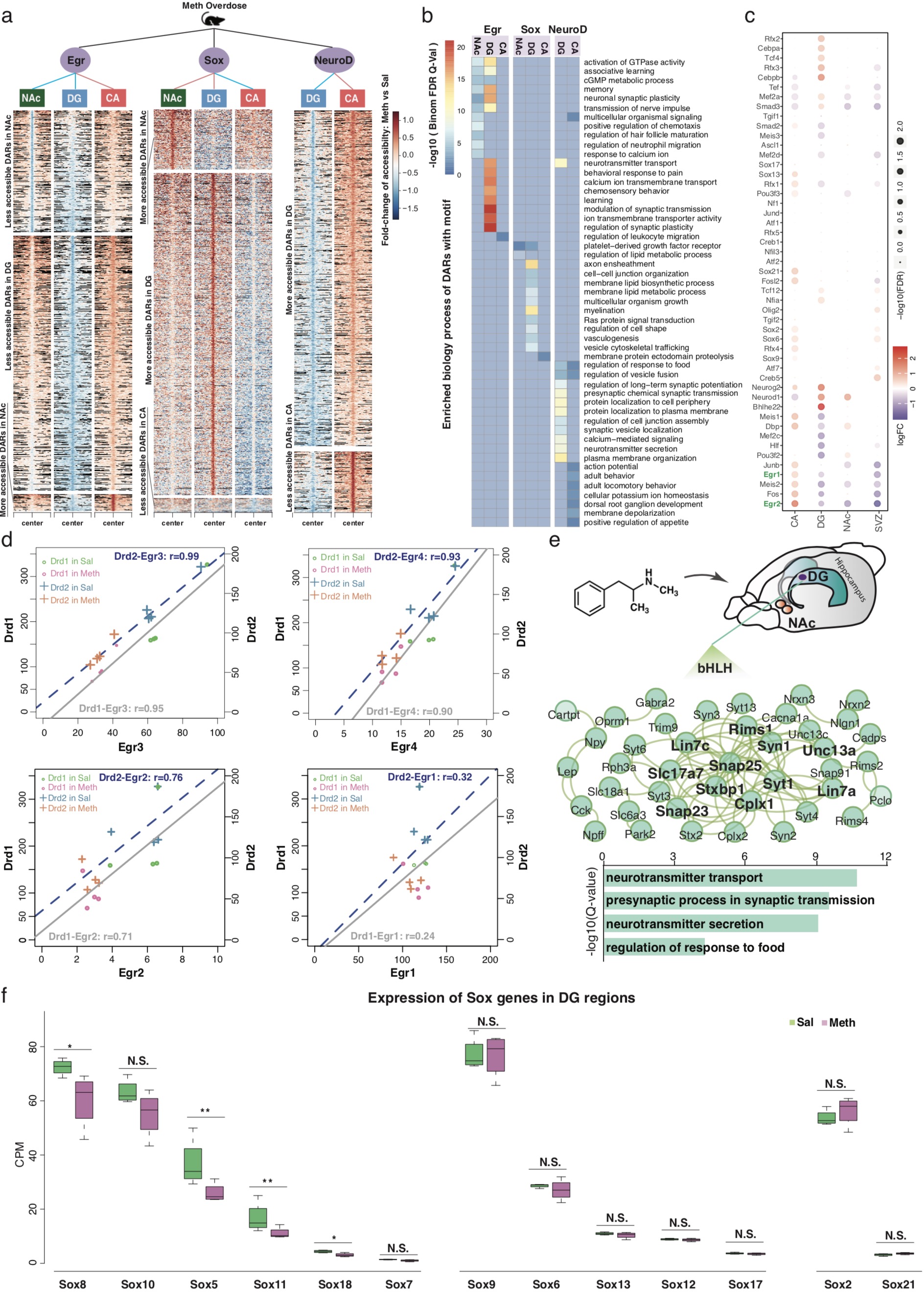

**Figure S5. METH-overdose induced DARs-associated regulatory networks in rat brain.** a, Heatmap of more and less accessible DARs contained the Egr, Sox and NeuroD binding motifs in different brain regions. b, Enriched biology processes based on DARs with Egr, Sox and NeuroD binding motifs. c, METH-overdose induced expression changes of transcription factors (TFs) whose binding motifs enriched in DARs. d, The correlation of expression changes between Drd1/2 genes and Egr family genes in rat NAc region with METH-overdose stimulus. e, Gene regulatory network and enriched biology process built by genes around DARs with bHLH binding motif in DG. f, Expression changes of Sox genes in DG region responding to METH-overdose.
